# Taxallnomy: Closing gaps in the NCBI Taxonomy

**DOI:** 10.1101/2020.05.28.119461

**Authors:** Tetsu Sakamoto, J. Miguel Ortega

## Abstract

NCBI Taxonomy is the main taxonomic source for several bioinformatics tools and databases since all organisms with sequence accessions deposited on INSDC are organized in its hierarchical structure. Despite the extensive use and application of this data source, taking advantage of its taxonomic tree could be challenging because (1) some taxonomic ranks are missing in some lineages and (2) some nodes in the tree do not have a taxonomic rank assigned (referred to as “no rank”). To address this issue, we developed an algorithm that takes the tree structure from NCBI Taxonomy and generates a hierarchically complete taxonomic tree. The procedures performed by the algorithm consist of attempting to assign a taxonomic rank to “no rank” nodes and of creating/deleting nodes throughout the tree. The algorithm also creates a name for the new nodes by borrowing the names from its ranked child or, if there is no child, from its ranked parent node. The new hierarchical structure was named taxallnomy and it contains 33 hierarchical levels corresponding to the 33 taxonomic ranks currently used in the NCBI Taxonomy database. From taxallnomy, users can obtain the complete taxonomic lineage with 33 nodes of all taxa available in the NCBI Taxonomy database. Taxallnomy is applicable to several bioinformatics analyses that depend on NCBI Taxonomy data. In this work, we demonstrate its applicability by embedding taxonomic information of a specified rank into a phylogenetic tree; and by making metagenomics profiles. Taxallnomy algorithm was written in PERL and all its resources are available at bioinfo.icb.ufmg.br/taxallnomy.

Database URL: http://bioinfo.icb.ufmg.br/taxallnomy

## INTRODUCTION

Any biological data are tightly linked to taxonomic data and several bioinformatics analyses depend on taxonomic information to achieve their objectives. Metagenomics, clinical forensic medicine, and other fields rely on fully-annotated taxonomic data to identify and group organisms present in a sample, often summarizing the results to a taxonomic rank such as family, order, class or phylum. Furthermore, any discussion made from evolutionary analyses refers to the taxonomic classification proposed so far. Taxonomic information can be obtained from several taxonomic databases, like the Catalogue of Life (1), which provides the taxonomic backbone to other projects such as Tree of Life (2), Encyclopedia of Life (3) and GBIF (4). Information provided by those databases is supported by taxonomy experts that feed other databases that cover a more specific clade, like FishBase (5), AmphibiaWeb (6), AnimalBase (7) and others. However, any analyses that involve molecular sequences are dependent on the NCBI Taxonomy (8), a reference taxonomic database with a huge compilation of taxonomic names and lineages of organisms that have a register of their DNA or protein sequence in one of the databases comprising the International Nucleotide Sequence Database Collaboration (INSDC) (9). Since INSDC comprises the three main molecular sequence repositories, GenBank, ENA and DBJJ, the information provided by NCBI Taxonomy is broadly used in biological databases covering diverse subjects that rely on data from INSDC, such as UniprotKB (10), Ensembl (11), Pfam (12), SMART (13), Panther (14), OMA (15) and miRBase (16). Moreover, other main primary biological databases, such as PDB (17), ArrayExpress (18) and KEGG (19) link their accessions to taxonomic data from NCBI Taxonomy database, demonstrating the undeniable contribution of this database to several bioinformatics fields.

The taxonomic classification comprising the NCBI Taxonomy follows the phylogenetic taxonomy scheme with the topology reflecting a consensus of views from taxonomic and molecular systematic literature (8). Each node of the tree represents a taxon, and each of them has a taxonomic name and a taxonomic identifier (txid). In addition, some nodes may have a taxonomic rank, which are similar to those used on the Linnaean classification system, such as Phylum, Class, and Order; and serve as important references of taxonomic classification. Several bioinformatics approaches rely on the rank-based classification provided by NCBI Taxonomy to make, for instance, taxonomic profiles of metagenomic data or to assist the taxonomic classification of sequence data. Besides the large use of rank information in the bioinformatics community, however, there are some important issues to be considered when managing these data. When querying for some organism lineages, we could observe that some of them are lacking some ranks. In a consultation on NCBI Taxonomy performed in May 2019, the lineage of the pig (*Sus scrofa*, NCBI:txid9823), for instance, had no taxon with the Order rank. Thale cress (*Arabidopsis thaliana*, NCBI:txid3702), on the other hand, did not contain a taxon with a Class rank in its lineage. If we look further in the taxonomic lineage, we could also find some taxa without a rank, denoted as “no rank” taxa. They add phylogenetic information to the taxonomy base, pointing out monophyletic groups.

These issues may be due to the uncertainty or conflict amongst experts on the classification of this group and turn to make hierarchical ranks of NCBI Taxonomy incomplete. Because of that, a simple query regarding the taxonomic ranks, such as “How many distinct taxa of class rank are represented in this data?” could become a difficult task. For instance, if the class for thale cress and several non-assigned classes of monocots are present they will all be counted as “NULL” in a computational database, grouping non-related counts. For such analyses, a hierarchically complete taxonomic tree incorporating taxonomic ranks could be of great benefit. Thus, in this work, we developed an algorithm that takes the taxonomic tree provided by NCBI Taxonomy and generates a hierarchical taxonomic tree in which all lineages have the same depth and all hierarchical levels corresponding to a taxonomic rank. The final database was named taxallnomy because it provides taxonomic names for all taxonomic ranks in a lineage comprising the tree. taxallnomy is in a tab-delimited format, making easy to access all members of a given clade. Users can access and explore the hierarchical structure of the taxallnomy database at its website at bioinfo.icb.ufmg.br/taxallnomy. Instructions to access the data programmatically through API or to produce the taxallnomy database in a local machine are also available at the taxallnomy website. Local production is very simple and grants the use of updated information.

## MATERIAL AND METHODS

### Dataset

Dump files with taxonomic information provided by NCBI Taxonomy FTP server (ftp.ncbi.nih.gov/pub/taxonomy/) were used for the construction of the taxallnomy database. The results presented in this work were obtained using dump files downloaded on May 16, 2019, although the website is kept up to date.

### Concepts

Here we present common terms used when referring to the hierarchical structure of NCBI Taxonomy. The taxonomic tree of NCBI Taxonomy consists of several **taxa** organized in a hierarchical data structure. All taxa have a name (e.g. *Homo sapiens*, Mammalia, Bacteria) and a numeric identifier (Taxonomy identifier or txid; e.g 9606 for *Homo sapiens*) associated to and they correspond to the nodes of the tree. Each taxon is connected to a single node of a level above (parent taxon), except for the **root node** which is positioned on the top of the tree. Furthermore, a taxon may be connected to one or more nodes of a level below (child taxon); when a taxon is not connected to any child taxon, it is referred to as **leaf taxon** (or leaf node). Each taxon also may have one of the 33 **taxonomic ranks** assigned (Table 1). Taxonomic ranks also follow a hierarchy such that a taxon of a higher rank cannot be a descendant of a taxon of a lower rank (e.g. a taxon of phylum rank cannot be a descendant of a taxon of class rank). In this work, we also refer to ranks through numbers which are the **rank levels**. The rank level ranges from 1 to 33, and the highest (Superkingdom) and the lowest (Forma) ranks have respectively the rank levels 1 and 33. Not all taxa on NCBI Taxonomy have a taxonomic rank assigned to itself and those taxa are referred to as “**no rank**” (e.g. Tetrapoda, NCBI:txid32523). A **taxonomic lineage** of a taxon is referred to as the route in the hierarchical structure which takes the taxon to the top of the hierarchy (in this case, the root node) and it could be composed of both taxonomically ranked and “no rank” nodes. Along the taxonomic lineage of a taxon, we could obtain the rank classification in each level and also verify that the lineages of some taxa may lack some taxonomic ranks (e.g. pig’s order and thale cress’s class). Finally, some nodes in the tree were considered as **unclassified taxa**, which include nodes that have in their lineage a taxon with the term unpublished, unidentified, unassigned, environmental samples, or *incertae sedis* on its name.

**Table 1:** Taxonomic ranks found in NCBI Taxonomy and abbreviations used

| rank level | rank name | abbrev. | rank priority | # of lineages<br>with the rank |
| --- | --- | --- | --- | --- |
| 1 | superkingdom | spKin | 1 | 1321086 |
| 2 | kingdom | Kin | 9 | 775938 |
| 3 | subkingdom | sbKin | 19 | 92281 |
| 4 | superphylum | spPhy | 32 | 164 |
| 5 | phylum | Phy | 6 | 1274596 |
| 6 | subphylum | sbPhy | 8 | 845043 |
| 7 | superclass | spCla | 20 | 86336 |
| 8 | class | Cla | 7 | 1137475 |
| 9 | subclass | sbCla | 10 | 579604 |
| 10 | infraclass | inCla | 14 | 381205 |
| 11 | cohort | Coh | 15 | 353345 |
| 12 | subcohort | sbCoh | 26 | 10665 |
| 13 | superorder | spOrd | 18 | 151859 |
| 14 | order | Ord | 5 | 1285474 |
| 15 | suborder | sbOrd | 11 | 450610 |
| 16 | infraorder | inOrd | 16 | 333382 |
| 17 | parvorder | prOrd | 21 | 61050 |
| 18 | superfamily | spFam | 13 | 409396 |
| 19 | family | Fam | 4 | 1318244 |
| 20 | subfamily | sbFam | 12 | 440875 |
| 21 | tribe | Tri | 17 | 216503 |
| 22 | subtribe | sbTri | 22 | 36549 |
| 23 | genus | Gen | 3 | 1319772 |
| 24 | subgenus | sbGen | 24 | 15337 |
| 25 | section | Sec | 28 | 4271 |
| 26 | subsection | sbSec | 31 | 303 |
| 27 | series | Ser | 33 | 155 |
| 28 | species group | Gro | 25 | 13963 |
| 29 | species subgroup | sbGro | 29 | 963 |
| 30 | species | Spe | 2 | 1319803 |
| 31 | subspecies | sbSpe | 23 | 27255 |
| 32 | varietas | Var | 27 | 7771 |
| 33 | forma | For | 30 | 486 |

### Database construction

The algorithmic challenge that we proposed for this work is to fill in the blanks on the taxonomic lineage considering the taxonomic ranks. To accomplish this, we created an algorithm that performs one of the two operations: (1) assigns taxonomic ranks to currently unranked taxa throughout the taxonomic tree if possible, or (2) creates ranked taxon nodes, naming it accordingly to its child or parent nodes. In both cases prefixes will distinguish taxallnomy additions from the original names, e.g. in Cla_eudicotyledons, the Class rank was assigned to the “no rank” node eudicotyledons (prefix Cla_) naming *A. thaliana* Class; in Order_of_Suina, the Order rank was assigned to a created node, parent of infraorder Suina (prefix Order_of_); and in SbSpe_in_Homo sapiens, the Subspecies rank was assigned to a created node, child of species *Homo sapiens*.

The algorithm begins evaluating all unranked nodes by moving through the hierarchical levels of the taxonomic tree, looking for unranked taxa. For each unranked taxon found, the algorithm evaluates if some of 33 taxonomic ranks occurring in NCBI taxonomy (Table 1) can be assigned to it. Since the taxonomic ranks follow a hierarchy, an unranked node can assume neither a rank level that is lower or equal than those found among its ascendant nodes nor a rank that is higher or equal than those found among its descendant nodes. So, to determine the ranks that an unranked node could be assigned with, the algorithm firstly verifies the highest and the lowest taxonomic rank levels found among its ascendant and descendant nodes, respectively. The rank levels that are between them are those that can be assigned to the unranked node in conformation with the rank hierarchy, thus considered as candidate ranks (numbers in balloons in Figure 1).

**Figure 1:**
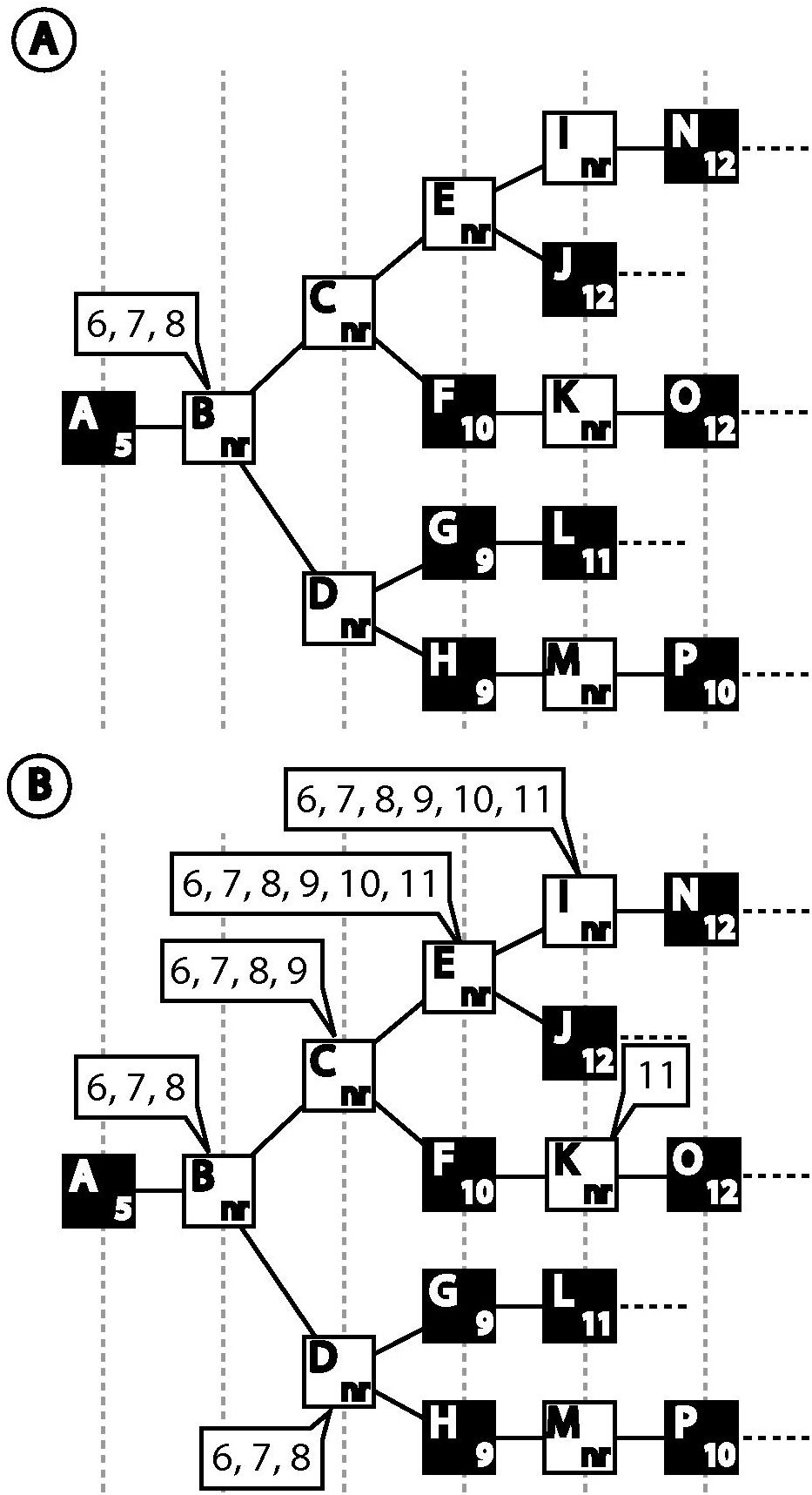
Evaluation of unranked nodes for rank assignment. Ranked nodes are denoted by filled squares. (A) In the hypothetical taxonomic tree, node B is between taxonomic rank level 5 (phylum) which is the lowest rank among its ascendant nodes; and taxonomic rank level 9 (subclass) which is the highest taxonomic rank among all its descendant nodes (9 to 12). Thus, the taxonomic ranks that node B could assume without affecting the rank hierarchy are subphylum (6), superclass (7) or class (8), in the balloon. (B) After evaluating all unranked nodes, we can find (i) those unranked nodes that cannot assume any ranks (node M, since it is between ranked nodes 9, subclass, and 10, infraclass) and (ii) those that could assume one or more ranks (nodes with a balloon).

After evaluating all unranked nodes, the algorithm proceeds to the rank assignment procedure. In this step, the algorithm goes through the taxonomic tree, starting from the root, looking for unranked nodes with candidate ranks to assign an appropriate rank to it. A simple case of this assignment occurs in nodes that have a single candidate rank without any unranked node as its parent or child (e.g. node K in Figure 1B). In this case, the algorithm simply assigns the candidate rank to the node. The assignment process becomes more complex when the unranked node has two or more candidate ranks and/or has additional unranked nodes among its child nodes since it enables more than a single valid way to perform the rank assignment. To deal with those situations, we created a set of algorithmic rules to decide the nodes and the taxonomic ranks to be used for the assignment (Figure 2A). The rules were designed aiming to assign ranks to as many unranked nodes as possible while prioritizing the assignment of those ranks most frequently found in the lineages of the taxonomic tree.

**Figure 2:**
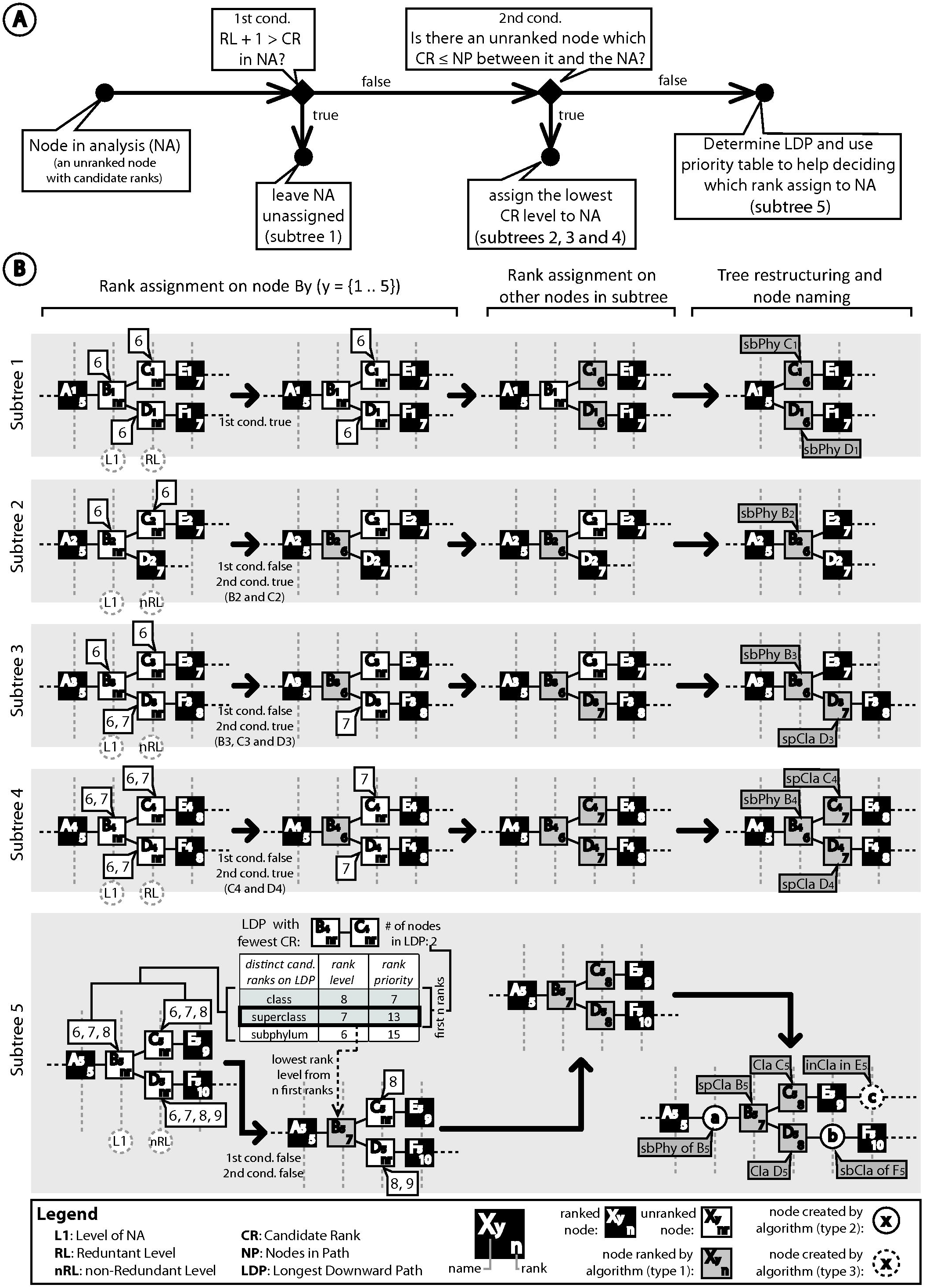
Rank assignment step. See text for a detailed explanation. (A) Set of rules followed by the algorithm to assign a single rank to an unranked node with candidate ranks. (B) Some examples of complex situations faced by the algorithm and the way it solves the rank assignment, the tree restructuring, and the node naming. Candidate ranks of an unranked node are in balloons. Square nodes represent taxa originally found in NCBI Taxonomy. Names assigned to nodes ranked (type 1) or created (types 2 and 3) by the algorithm are shown in gray balloons.

For a better understanding of the assignment rules, consider the subtrees in Figure 2B which illustrates different situations found by the algorithm for the rank assignment problem. In all subtrees, the node in analysis (NA) is the unranked node B_n_ (n = {1, 2, …, 5}). Also, the hierarchical level of the NA is referred to as the first level (L1). The first condition evaluated by the algorithm is the existence and the number of levels that are redundant with the L1 (RL). We consider that levels following the L1 are redundant if all nodes on it are (1) unranked and (2) have the same candidate ranks as the NA, and if (3) the level above it is the L1 or a redundant level without leaf nodes. If the number of candidate ranks in the NA is less than the number of levels that needs a rank (L1 plus consecutive redundant levels), there are not enough ranks to assign to the nodes on those levels. In this case, one option is to assign the lowest rank level among the candidate ranks to the NA and leave some of its descendant nodes unranked. However, we opted to leave the NA without a rank so that the unranked nodes of further levels could have a rank assigned. This procedure was chosen as this could result in more unranked nodes with a rank assigned, favoring the dichotomization of the final tree. In subtree 1 (Figure 2B), the NA (B_1_) has one candidate rank (taxonomic level 6), and the level following the L1, which is composed of nodes C_1_ and D_1_, meets all conditions established to be a redundant level (RL). Since the number of candidate ranks in the NA (CR = 1) is not sufficient to rank the nodes on L1 and on further redundant level (L1+RL = 2), the algorithm leaves the node B_1_ without a rank, allowing the nodes of further level (C_1_ and D_1_) to have a rank assigned.

If the previous condition is not true then the next condition evaluated by the algorithm is the existence of unranked nodes in the subtree in which the number of candidate ranks on it is equal to or less than the number of consecutive unranked nodes in the path between it and the NA. If a node in this condition is found (Figure 2B, subtrees 2, 3 and 4), it is likely that all candidate ranks on it could be distributed along with the unranked nodes in the path linking it to the NA. So, in this condition, the algorithm assigns the lowest rank level among the candidate ranks to the NA.

If none of those conditions apply, it is an indicator that the subtree cannot bear to assign all candidate ranks to its nodes (Figure 2B, subtree 5). In this case, the algorithm has to decide the ranks to be used for the assignment process. To help with this, we determined an order of priority of which taxonomic ranks should be firstly assigned based on the frequency that they appear in the leaf lineages (Table 1). The more frequent the rank, the higher is its priority. To make use of this order of priority, the algorithm searches for the longest downward path (LDP) of consecutive unranked nodes, starting from the NA. Once the LDP is found, the algorithm stores the number of nodes on this path and the distinct candidate ranks found among the nodes comprising the path. If there is more than one LDP in the subtree, the algorithm considers the one with less distinct candidate ranks along the path. Then, the candidate ranks are sorted according to their order of priority and the first *n* ranks, in which *n* is the number of nodes in the LDP, are extracted. The extracted ranks will be those to be assigned to the nodes in the LDP. Since the NA is the first node in the LDP, the algorithm picks the rank of lowest level among the extracted ranks and assigns it to the NA.

After an unranked node has a rank assigned, the further unranked nodes have their list of candidate ranks updated and visited by the algorithm to perform the same analysis. After performing this procedure on all unranked nodes, all of them will have a single rank or no rank assigned (Figure 2B).

The final step of the algorithm consists of creating and deleting nodes to make the taxonomic tree complete hierarchically; and on defining a name for the created nodes and for the unranked taxa which had a rank assigned (Figure 2B, tree restructuring and node naming step). For this, the algorithm will delete all unranked taxa that did not have a rank assigned in the previous procedures; as occurred with the nodes “B_1_”, “C_2_” and “C_3_” (Figure 2B, subtrees 1, 2 and 3). On the other hand, unranked taxa with a rank assigned are maintained and new names are assigned to them to indicate that they were originally unranked. The new names of these nodes consist on the abbreviation of the rank assigned (Table 1) followed by the original name of the node; i.e. in the node “C_1_” (Figure 2B, subtree 1), since it has the Subphylum rank assigned, its new name will be “sbPhy C_1_”. The nodes that meet this condition are referred to as taxa of **type 1**. An example of a node of this type in the taxallnomy tree is cla_eudicotiledons, which is the proposal for thale cress’ class.

The taxonomic tree has portions where two consecutive taxa do not have consecutive ranks. In this case, the algorithm creates nodes between them and assigns the created nodes with ranks that are missing. For instance, we could observe in subtree 5 of Figure 2B that there should be nodes with ranks of Subphylum (level 6) between the nodes “A_5_” (Phylum, level 5) and “B_5_” (Superclass, level 7). To fulfill this gap, the algorithm creates between them a node (node “a”) with Subphylum rank assigned to it. This type of node is referred to as **type 2** and is named using the abbreviation of the assigned rank followed by the preposition “of” and the original name of its first ranked descendant node. For the node “a”, since it has the node “B_5_” as the first ranked descendant node, it is named as “sbPhy of B_5_”. Pig’s order, for example, is proposed to be ord_of_Suina, which Suina is an Infraorder stated in the original database.

Finally, if there are some lineages with missing ranks because there is no node of a higher level, the algorithm will also visit these lineages and create a node for each missing rank. In subtree 5 of Figure 2B, the node “E_5_” is a leaf node of Subclass rank (level 9). Since “E_5_” is a leaf node, all ranks after Subclass are missing in this lineage. In this case, taxallnomy will visit these nodes and create nodes to fulfill those missing ranks. The node “c” in subtree 5 (Figure 2B) is a node created for this proposal. To name this node, the algorithm takes the abbreviation of the missing rank followed by the preposition “in” and by the original name of the last taxon of the lineage (“ifc in E_5_”). These nodes are referred to as taxa of **type 3**. They are useful in cases when there are, for example, subspecies declared in the database. For instance, *Sus scrofa* (NCBI:txid9823) has over 60 thousand proteins deposited, but only around 1.5 thousand are assigned to one of its 11 subspecies. Therefore, most of those entries are “NULL” for the Subspecies rank in the original database; but, by creating the node of type 3, all of them are treated to have a node of Subspecies rank named “sbSpe in Sus scrofa”. Another usage of nodes of type 3 is on metagenomics analysis (Figure 7), when there are entries annotated with taxa of lower rank levels and one wants to count the number of distinct taxa of higher rank levels.

Species and Genus are ranks with high frequency (Table 2), thus both have high priority during the rank assignment procedure. Therefore, an unranked node that has one of those ranks as candidates are more likely to have one of them assigned. We evaluated some lineages of leaf taxa lacking for Species or Genus ranks and verified that some unranked nodes are appropriate to have one of those ranks. For instance, in an older version of NCBI Taxonomy (September 19, 2016), *Beringia wynnei* (NCBI:txid1037071) was a leaf taxon of Species rank that did not have a Genus rank in its lineage. However, its lineage contained an unranked node named *Beringia* (NCBI:txid1037069), which had the Genus rank appropriately assigned by the algorithm. Similarly, *Nocardia argentinensis* ATCC 31306 (NCBI:txid1311813), in the same version of NCBI Taxonomy, was a “no rank” leaf taxon, which did not have a node with Species rank in its lineage, but it contained an unranked node named *Nocardia argentinensis* (NCBI:txid1311812). The current algorithm also appropriately assigned the Species and Subspecies ranks to the nodes *N. argentinensis* and *N. argentinensis* ATCC 31306, respectively. However, depending solely on these rules incurs some obvious errors, like in those unranked leaf taxa which do not have nodes with Species and Genus rank in its lineage. For instance, Rosodae (NCBI:txid721787) is a “no rank” leaf taxon that has a parent node with Subfamily rank (level 20). According to the algorithm, Rosodae could have ranks ranging from Tribe (level 21) to Forma (level 33), and, based on the rank priority, it would be assigned to Species rank, which is not a proper rank for it. To correct this situation, special rules were added to the algorithm to have the Species and Genus ranks assigned to an unranked node. We established that the Species rank assignment to an unranked node should occur only if, among its ascendant nodes, there is a node of Genus rank in the original database. On the other hand, an unranked node should have the Genus rank assigned if there are nodes of Species rank among its descendants in the original database. Moreover, the assignment of both ranks to an unranked node should not occur if a node has terms in its name that identify it as an unclassified entry. With these rules, the unranked node Rosodae mentioned before has the Tribe rank assigned instead of Species rank.

**Table 2:** BLAST result with taxonomic data from taxallnomy

| Entry | E-value | Ident | Taxid | Class<sup>1</sup> | Order<sup>2</sup> | Species |
| --- | --- | --- | --- | --- | --- | --- |
| P04637 | 0 | 1 | 9606 | Mammalia | Primates | *Homo sapiens* |
| H2QC53 | 0 | 1 | 9598 | Mammalia | Primates | *Pan troglodytes* |
| G3R2U9 | 0 | 0,997 | 9595 | Mammalia | Primates | *Gorilla gorilla* |
| H2NSL2 | 0 | 0,977 | 9601 | Mammalia | Primates | *Pongo abelii* |
| A0A0D9RG12 | 0 | 0,959 | 60711 | Mammalia | Primates | *Chlorocebus sabaeus* |
| P56424 | 0 | 0,957 | 9544 | Mammalia | Primates | *Macaca mulatta* |
| G7PTI9 | 0 | 0,957 | 9541 | Mammalia | Primates | *Macaca fascicularis* |
| A0A096P2R8 | 0 | 0,957 | 9555 | Mammalia | Primates | *Papio anubis* |
| G1RF61 | 0 | 0,957 | 61853 | Mammalia | Primates | *Nomascus leucogenys* |
| F7GNX0 | 0 | 0,896 | 9483 | Mammalia | Primates | *Callithrix jacchus* |
| I3N5N2 | 0 | 0,865 | 43179 | Mammalia | Rodentia | *Ictidomys tridecemlineatus* |
| Q95330 | 0 | 0,86 | 9986 | Mammalia | Lagomorpha | *Oryctolagus cuniculus* |
| G5B5D6 | 0 | 0,832 | 10181 | Mammalia | Rodentia | *Heterocephalus glaber* |
| A0A091CJU7 | 0 | 0,827 | 885580 | Mammalia | Rodentia | *Fukomys damarensis* |
| H0XGB0 | 0 | 0,833 | 30611 | Mammalia | Primates | *Otolemur garnettii* |
| T0MFN1 | 0 | 0,809 | 419612 | Mammalia | Ord_of_Tylopoda | *Camelus ferus* |
| G1MEP6 | 0 | 0,807 | 9646 | Mammalia | Carnivora | *Ailuropoda melanoleuca* |
| Q9TUB2 | 0 | 0,825 | 9823 | Mammalia | Ord_of_Suina | *Sus scrofa* |
| F1PI27 | 0 | 0,807 | 9615 | Mammalia | Carnivora | *Canis lupus* |
| P41685 | 0 | 0,805 | 9685 | Mammalia | Carnivora | *Felis catus* |
| P67939 | 0 | 0,802 | 9913 | Mammalia | Ord_of_Ruminantia | *Bos taurus* |
| C3VC56 | 0 | 0,792 | 9940 | Mammalia | Ord_of_Ruminantia | *Ovis aries* |
| Q9WUR6 | 0 | 0,782 | 10141 | Mammalia | Rodentia | *Cavia porcellus* |
| M3YC88 | 0 | 0,789 | 9669 | Mammalia | Carnivora | *Mustela putorius* |
| L5JZ91 | 0 | 0,774 | 9402 | Mammalia | Chiroptera | *Pteropus alecto* |
| P10361 | 0 | 0,77 | 10116 | Mammalia | Rodentia | *Rattus norvegicus* |
| P02340 | 0 | 0,774 | 10090 | Mammalia | Rodentia | *Mus musculus* |
| G3T035 | 0 | 0,773 | 9785 | Mammalia | Proboscidea | *Loxodonta africana* |
| L9KM90 | 0 | 0,816 | 246437 | Mammalia | Scandentia | *Tupaia chinensis* |
| L7N1B2 | 0 | 0,741 | 59463 | Mammalia | Chiroptera | *Myotis lucifugus* |
| L5M0C7 | 0 | 0,727 | 225400 | Mammalia | Chiroptera | *Myotis davidii* |
| S7NG87 | 0 | 0,752 | 109478 | Mammalia | Chiroptera | *Myotis brandtii* |
| F6TL72 | 2,00E-173 | 0,9 | 9796 | Mammalia | Perissodactyla | *Equus caballus* |
| G3WS63 | 3,00E-169 | 0,653 | 9305 | Mammalia | Dasyuromorphia | *Sarcophilus harrisii* |
| H3B1Z4 | 3,00E-128 | 0,53 | 7897 | Cla_Coelacanthimorpha | Coelacanthiformes | *Latimeria chalumnae* |
| P10360 | 5,00E-127 | 0,532 | 9031 | Aves | Galliformes | *Gallus gallus* |
| A0A151MW63 | 2,00E-124 | 0,535 | 8496 | Cla_of_Crocodylia | Crocodylia | *Alligator mississippiensis* |
| W5N8V9 | 7,00E-120 | 0,505 | 7918 | Actinopteri | Semionotiformes | *Lepisosteus oculatus* |
| K7G3P4 | 7,00E-117 | 0,506 | 13735 | Cla_of_Testudines | Testudines | *Pelodiscus sinensis* |
| F7A9M9 | 1,00E-111 | 0,491 | 8364 | Amphibia | Anura | *Xenopus tropicalis* |
| P79734 | 2,00E-110 | 0,575 | 7955 | Actinopteri | Cypriniformes | *Danio rerio* |
| M7CJF0 | 6,00E-108 | 0,585 | 8469 | Cla_of_Testudines | Testudines | *Chelonia mydas* |

<sup>1</sup> taxa of Class rank created by taxallnomy begin with "cla\_".<sup>2</sup> taxa of Order rank created by taxallnomy begin with "ord\_".

### Taxonomic code for taxallnomy taxa

The primary identification of each node comprising the taxallnomy tree is the Taxonomy ID provided by the NCBI Taxonomy database. However, since the taxallnomy algorithm assigns ranks to nodes and creates new nodes, we formulated a code that properly identifies them. The taxallnomy code consists of three digits added as a decimal number in the Taxonomy ID of each node. The first two digits indicate the taxonomic rank in which it was assigned. It goes through the code “01” to “33”, in which the first code (“01”) refers to the Superkingdom rank and the last one (“33”) refers to the Forma rank. The third digit ranges from 1 to 3 and indicates the approach used by the algorithm to create/modify a node. The codes 1, 2 and 3 refer respectively to taxa of type 1, type 2 and type 3. For instance, in the taxon code 6072.031, 6072 corresponds to the NCBI Taxonomy ID (Eumetazoa) and 031 is the code added by the taxallnomy algorithm, indicating that it is a node of type 1 created on Subkingdom rank. Using the taxallnomy name convention, the name of this node will be “sbKin Eumetazoa”. Furthermore, taxa originally ranked in the NCBI Taxonomy database has the code 000 included (e.g. 9606.000, that stands for the species *Homo sapiens*).

### Taxallnomy usability and availability

Users can query the taxallnomy database and download the results using its web interface at bioinfo.icb.ufmg.br/taxallnomy. In the web interface, users can also find an interactive taxallnomy tree, which allows easy exploration of its hierarchical structure. Advanced users can also programmatically query taxallnomy database using our REST service for this database (see taxallnomy web page for more instructions).

Users with high demand can also find all necessary files to have a copy of the taxallnomy database in a local MySQL database at the taxallnomy SourceForge page (sourceforge.net/projects/taxallnomy). Taxallnomy database is comprised of five main tables named “lin”, “lin_name”, “tree_complete”, “tax_data” and “rank”. The first two tables have the taxonomic lineages that comprise the taxallnomy tree. The tables have a column containing the NCBI Taxonomy ID (txid), which is the primary key column of the tables; and 33 columns representing the 33 taxonomic ranks found in the NCBI Taxonomy database. In the “lin” table, the taxonomic rank columns are filled with taxonomic codes, whereas, in “lin_name, those columns are filled with taxonomic names. The table “tree_complete” contains all parent-child relationships in the taxallnomy database that make the hierarchical structure complete. Two other hierarchically incomplete versions of the tree table are also available in the taxallnomy data source; one is the table “tree_all”, which includes the unranked nodes that did not have a rank assigned, and the other is the table “tree_original”, which has the same hierarchical structure as the one provided by the NCBI Taxonomy database. In “tax_data” table, users can find information about each taxon comprising the tree, such as its scientific name, common name and rank level. Finally, the “rank” table contains information about the ranks comprising the taxonomic tree, such as name, level, priority order, and abbreviation.

Since the NCBI Taxonomy database is frequently updated, the database used in the taxallnomy web page and provided in its SourceForge page is updated weekly. Users with a local copy of the taxallnomy database can acquire the updated database from its SourceForge page. Alternatively, we also provide a Perl script with the taxallnomy algorithm implemented at https://github.com/tetsufmbio/taxallnomy. The script can be executed in a UNIX system with an internet connection, which is required for downloading the latest version of the NCBI Taxonomy database.

## RESULTS

The taxonomic tree currently provided by NCBI Taxonomy has two issues regarding the taxonomic ranks: (1) some ranks are absent and (2) some taxa do not have a rank in a taxonomic lineage. The irregularity of its hierarchical structure is exemplified in Figure 3 (upper tree) which we take a portion of the taxonomic tree comprising some Classes of Kingdom Metazoa. In this context, we proposed to develop an algorithm that takes the tree structure from NCBI Taxonomy and make it complete regarding the taxonomic ranks. The new taxonomic database was named taxallnomy since it provides names for all ranks that are missing in a taxonomic lineage. By taking the equivalent portion of the tree from the taxallnomy database (Figure 3, lower tree) we could observe that all taxa with the same rank are positioned in the same hierarchical level. To achieve this, taxallnomy algorithm assigned ranks to some unranked taxa, such as Eumetazoa (level 3), Deuterostomia (level 4) and Panarthropoda (level 4); deleted others, such as Bilateria, Vertebrata and Gnathostomata; and created nodes to fill the missing ranks in a lineage, such as “spPhy (Superphylum) of Cnidaria”, “sbPhy (Subphylum) of Hexapoda”, “spCla (Superclass) of Chondrichthyes” and others.

**Figure 3:**
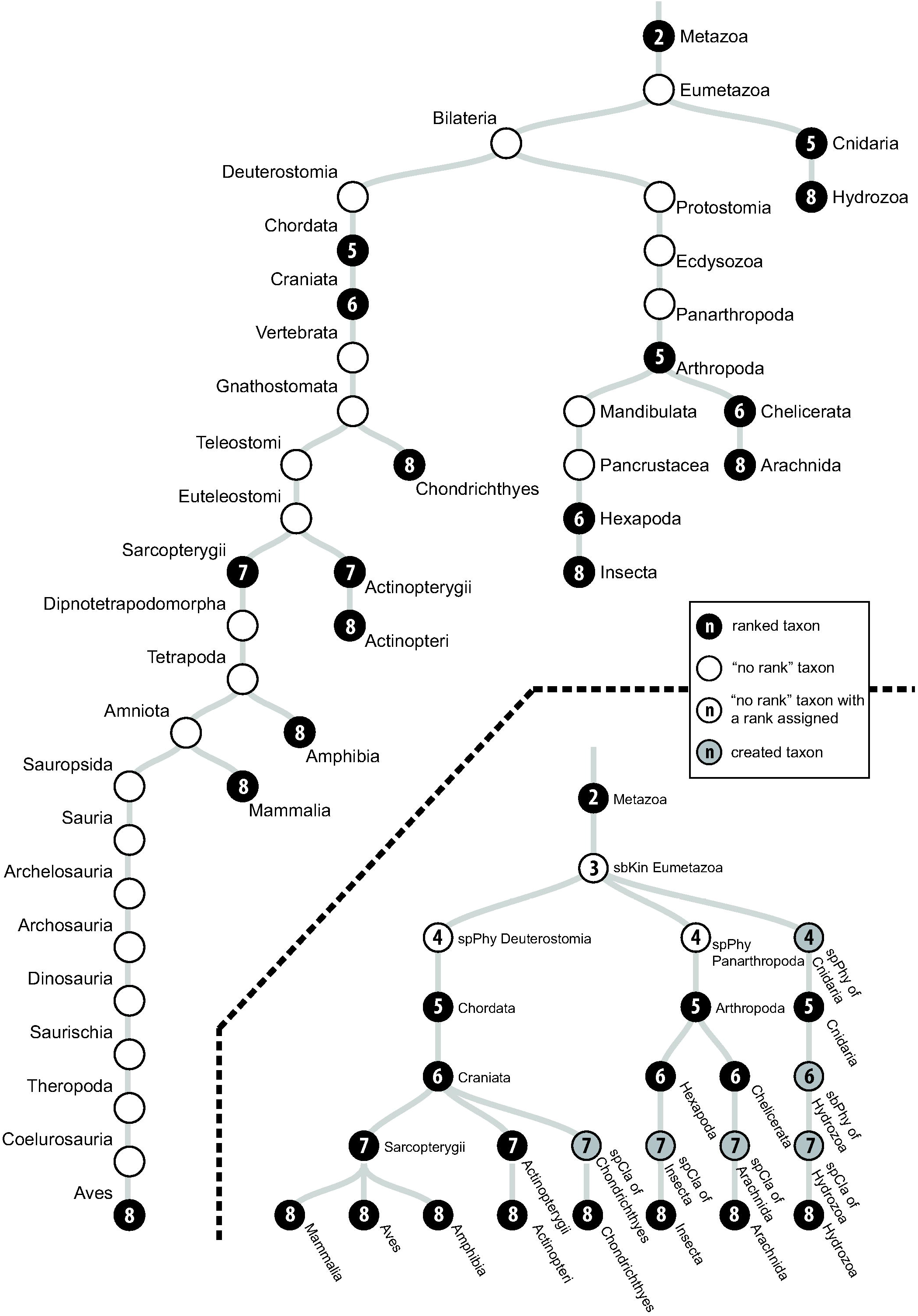
Portion of the taxonomic tree from NCBI Taxonomy (upper tree) and taxallnomy (lower tree) comprising some classes of Kingdom Metazoa. The number in the node (n) indicates the taxonomic rank level (see Table 1).

Observing the exemplified portion of the taxallnomy tree in more detail, one could question why the algorithm ranked the taxa Deuterostomia and Panarthropoda to Superphylum (level 4) instead of ranking the taxon Bilateria. Another questionable point can be found in the lineage of Insecta in which the algorithm did not rank the taxa Mandibulata or Pancrustacea to Subphylum (level 6), but created a new node (sbPhy of Hexapoda) instead. All those ranking patterns executed by the algorithm were established in order to be in agreement with the rank hierarchy. The taxon Bilateria could not have the rank Superphylum assigned because one of its descendant taxa (Scalidophora – not shown) has this rank. In a similar way, Mandibulata and Pancrustacea could not have the rank Subphylum assigned because both taxa have descendant taxa (e.g. Crustacea - not shown) with this rank.

Taxallnomy database consists of a total of 11,812,306, in which 10,573,069 (89.51%) of them are nodes created by the taxallnomy algorithm or unranked taxa that had a rank assigned. Among them, 214,503 (2.03%) are of type 1, 6,808,044 (64.39%) of type 2 and 3,550,522 (33.58%) of type 3. Moreover, the number of unranked nodes used to create the nodes of type 1 corresponds to 99.88% of all unranked nodes found in the original tree (214,754 nodes). The number of leaf taxa totalized 1,321,086, in which 60.34% are from Eukaryota, 26.51% from Bacteria, 0.37% from Archaea and 12.76% from Virus or Viroids Superkingdoms. In these countings, unclassified taxa (including unpublished, unidentified, unassigned, environmental, or *incertae sedis* taxa) were not included.

Since the taxallnomy tree is hierarchically complete and consequently all taxonomic lineages have all nodes of each rank level, the number of distinct taxa found in each rank expectedly increases as we go through the ranks (Figure 4). This contrasts with the original tree from the NCBI Taxonomy database, which shows a wide fluctuation in the number of distinct taxa along with the ranks. The contribution of the taxallnomy database in creating nodes and names can be noticed by measuring the differences in the number of distinct taxa on both trees.

**Figure 4:**
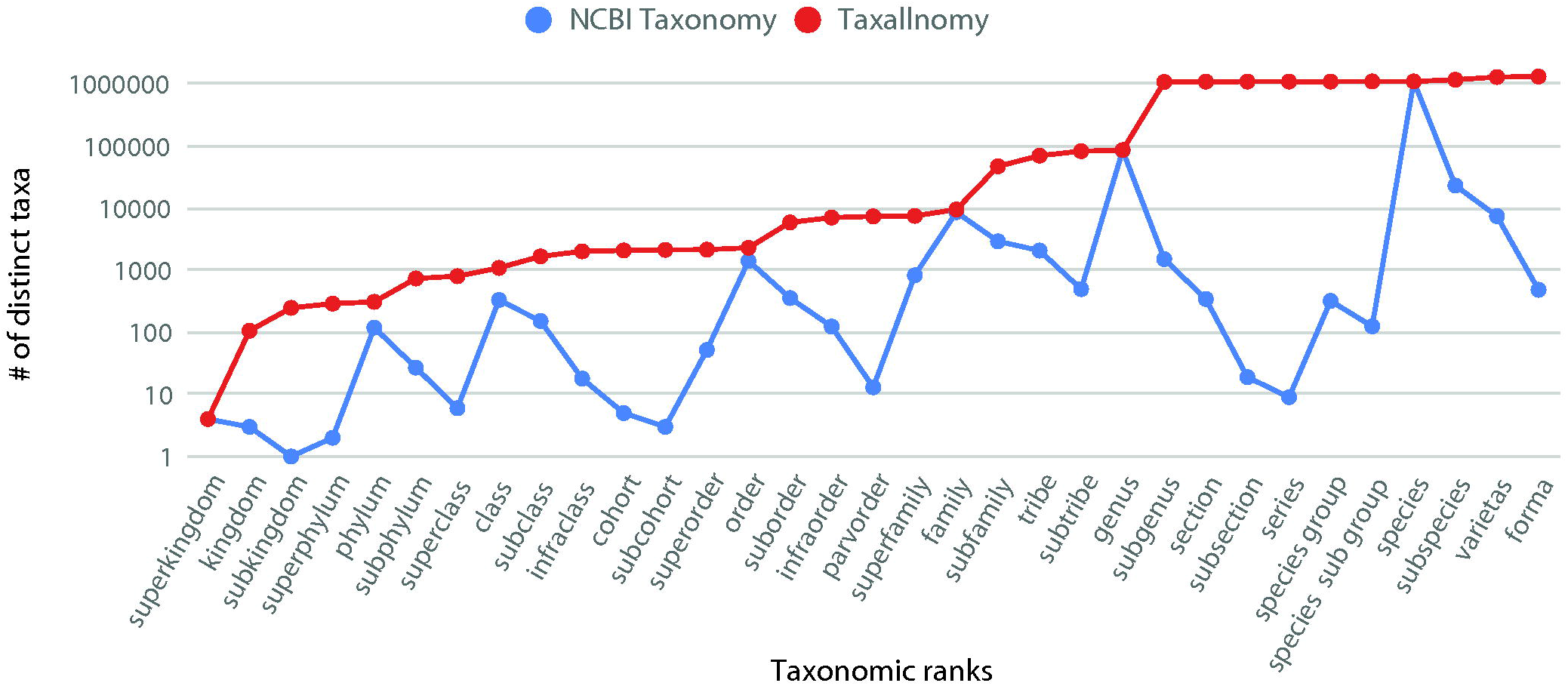
Number of distinct taxa in each taxonomic ranks in NCBI Taxonomy and taxallnomy databases.

Considering the contribution of taxallnomy in the taxonomic lineages of leaf taxa (Figure 5), we could firstly observe that some ranks are found in almost all lineages in the original tree. These ranks are referred here as **main** ranks and include Superkingdom, Phylum, Class, Order, Family, Genus, and Species. The other ranks, in contrast, are found originally in few lineages and had a node included in their lineages by the taxallnomy algorithm. Nodes of type 1 are found mainly in the first ranks (from Kingdom to Class ranks) and in Subspecies ranks, indicating the existence of unranked taxa in this range on the original tree worthy to be ranked. We also highlight that, despite Phylum and Class are part of main ranks, the taxallnomy algorithm has fulfilled those ranks in a considerable amount of lineages with taxa of type 1. Taxonomic lineages exhibit a great amount of type 2 nodes on rank levels lower than Species rank level and that are not part of main ranks. This occurs because there are no or few unranked taxa to assign a rank in those ranges, which forces the algorithm to create new nodes in the original tree. Finally, the nodes of type 3 are concentrated in the last ranks (from Subspecies to Forma ranks). This indicates that many leaf taxa analyzed are from Species rank, causing the algorithm to create taxa of type 3 for the further ranks.

**Figure 5:**
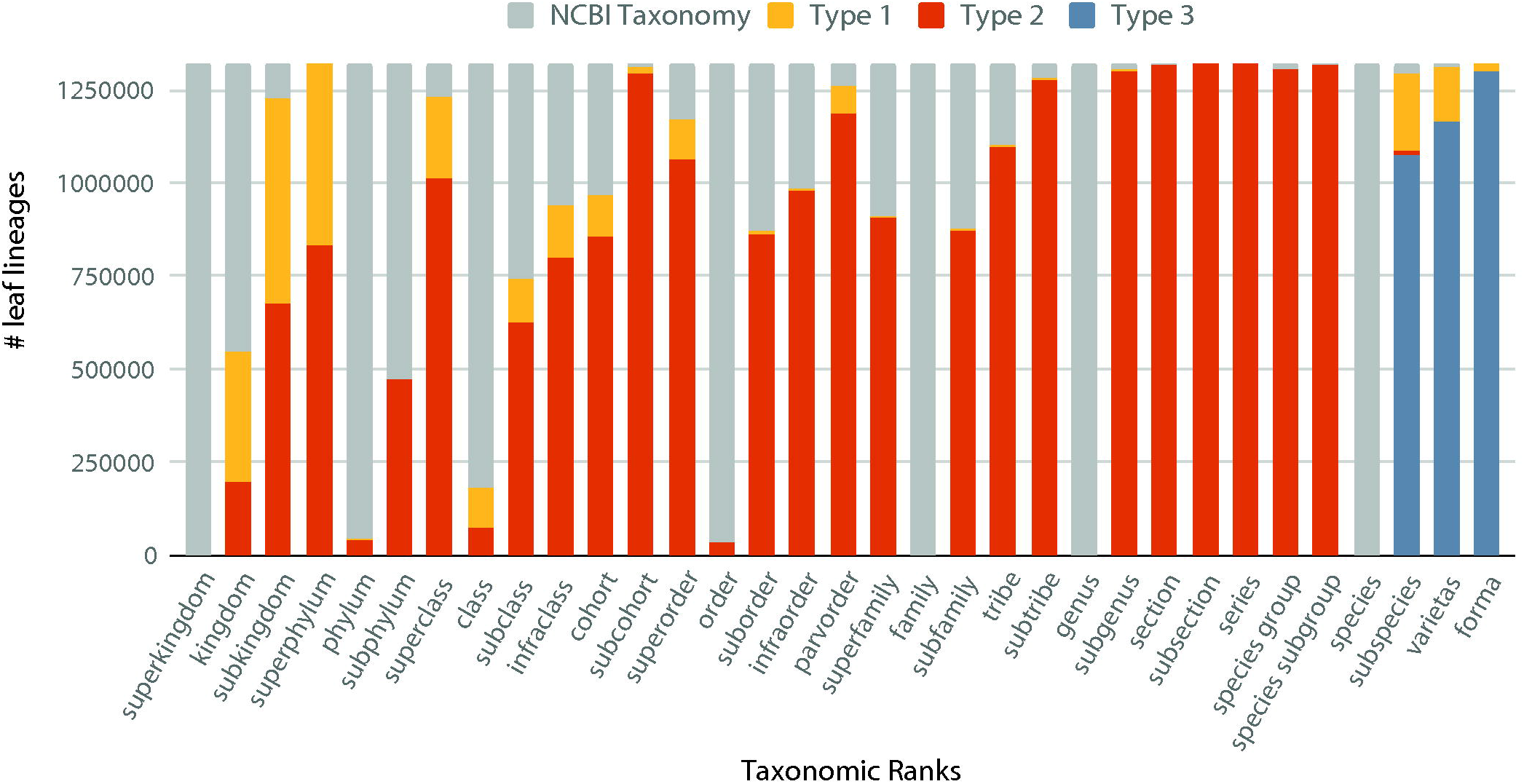
Frequency of taxa originally ranked in NCBI Taxonomy and taxa created by the taxallnomy algorithm (type 1, type 2 or type 3) in each taxonomic rank. A total of 1,321,086 taxonomic lineages of leaf taxa were evaluated.

## TAXALLNOMY APPLICABILITY

The lack of a taxon on a specified rank in a lineage could be inconvenient for any analysis in which we ask something about the taxonomic ranks on our data. One could take a simple BLAST result and ask which taxa from a specified rank are found among the subjects retrieved. If one tries to answer this using the original data from the NCBI Taxonomy database, he could come across subjects belonging to species that do not have a taxon with the queried rank. In this situation, we could take advantage of the taxallnomy database, which has the gaps of all taxonomic lineage fulfilled. For instance, taking a BLAST (20) result that used the Human P53 protein as query against the UniProt database (10) (Table 2), we could observe that most of the subjects retrieved in this analysis belongs to organisms that have a taxon with Class rank (Mammalia, Aves, Amphibia, and Actinopteri) in the original database, but some of them have a Class rank created by taxallnomy (“Cla_Coelacanthimorpha”, “Cla_of_Crocodylia” and “Cla_of_Testudines”). Without this information, we could not have an idea if those subjects are from organisms of the same Class or not. If we consider now the Order rank, we could observe that there are four subjects belonging to organisms that lack this rank in their lineage. By fulfilling those spots with information from taxallnomy, we have the four organisms classified in three distinct Orders (“Ord_of_Tylopoda”, “Ord_of_Suina” and “Ord_of_Ruminantia”).

In a similar way, taxonomic data are frequently incorporated into a phylogenetic tree to evidence some taxonomic groups. By embedding taxonomic data from taxallnomy to a phylogenetic tree, a user can select a rank and evidence taxa comprising the selected rank without worrying about the missing ranks. We exemplify this by evidencing the distinct Orders comprising a phylogenetic tree generated using the tumor protein 53 (P53) sequences of species from Superorder Laurasiatheria (Figure 6). In this tree, we could evidence three taxa that were created by taxallnomy: Ord_of_Ruminantia, Ord_of_Suina, and Ord_of_Tylopoda.

**Figure 6:**
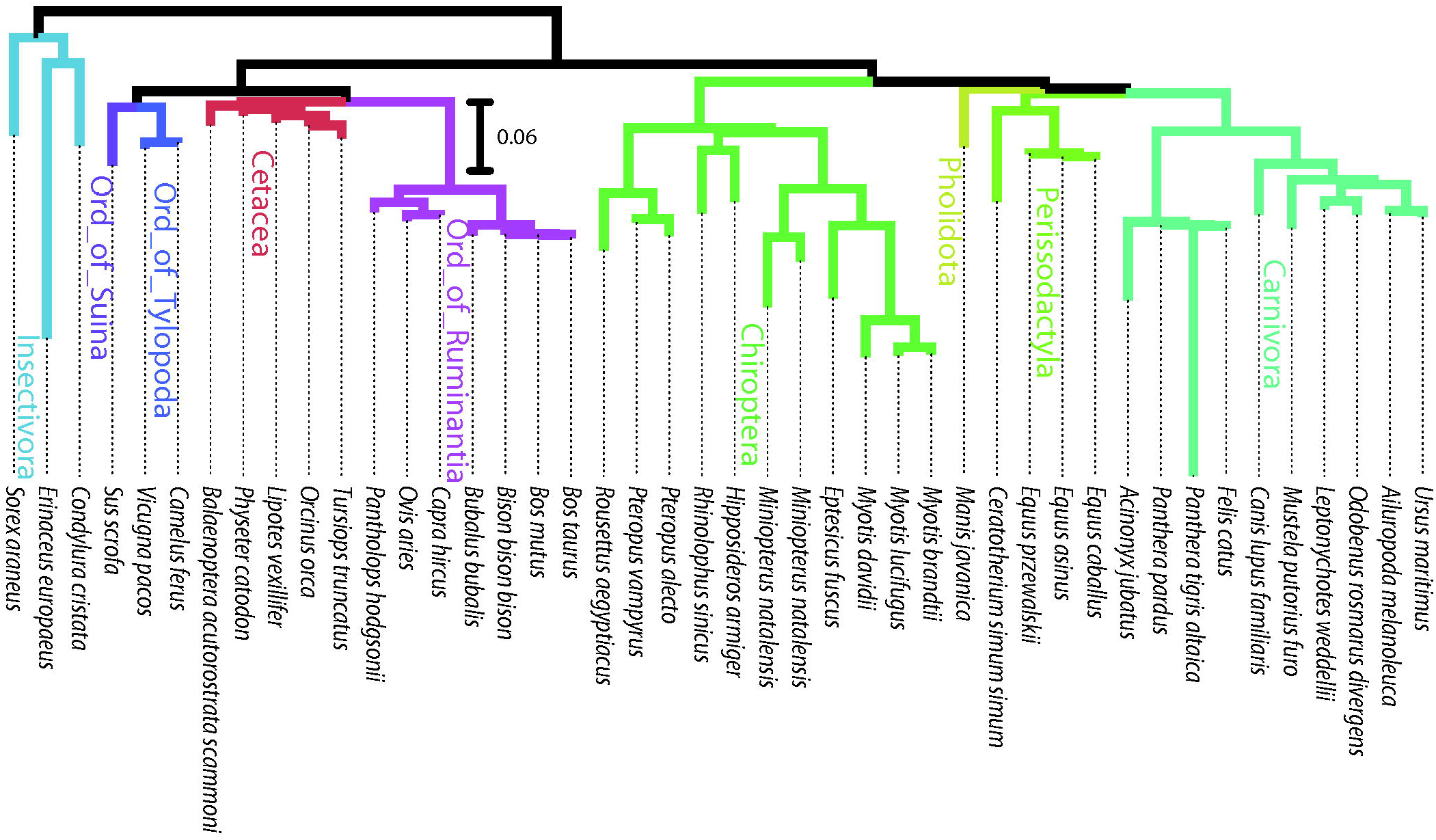
Phylogenetic tree of tumor protein P53 of species from Superorder Laurasiatheria. Branch color evidences the distinct taxa of Order rank comprising the tree. Orders beginning with “Ord_of_” (in bold) are taxa created by the taxallnomy algorithm.

Metagenomics analyses heavily rely on taxonomic data and information about taxonomic ranks. After the taxonomic annotation performed by software like MEGAN (21), MG-RAST (22) or the pipeline from EBI Metagenomics (23), researchers in this field seek a metagenomics profile to verify which taxa are predominant in an environmental sample. Since taxonomic annotation performed by those programs is based on the NCBI Taxonomy database, the taxonomic profile is usually performed by firstly extracting the taxonomic lineages of those taxa that were assigned to a read and then plotting the abundance of taxa in each taxonomic rank separately. However, as stated initially, some ranks are missing in some taxa, which oblige us, in the end, to include all those taxa without rank in a separate group or to omit them in the graphic representation. The same procedures are taken in the case in which there is a read annotated to a taxon of lower rank level and we want to have a taxonomic profile of a higher rank level. An alternative representation of the taxonomic profile is to show the taxa abundance along the taxonomic tree without accounting for the taxonomic rank. The problem of this approach is the occurrence of different taxonomic ranks in the same level of the tree since the depth of taxonomic lineages could vary between taxa. All these issues can be resolved by using a hierarchically complete taxonomic tree provided by the taxallnomy database. To exemplify this, we took a metagenomics sample collected from a tropical freshwater reservoir in Brazil (projectID on MG-RAST:mgp13799) and generated its taxonomic profiles using taxonomic sources from NCBI Taxonomy and taxallnomy. For this, we submitted the reads to MEGAN for taxonomic annotation and retrieved the taxonomic lineage from both databases for each taxon that appeared in the annotation process. Then, we assembled all taxonomic lineages retrieved in a spreadsheet and generated, for instance, pie charts for the ranks Kingdom, Phylum, and Class (Figure 7). In the profile obtained using NCBI Taxonomy (Figure 7A), depending on the metagenomic sample and taxonomic rank in analysis, several reads would be omitted or grouped in the unclassified group since the lineage of the taxa assigned to them miss for those ranks. Since lineages retrieved from the taxallnomy database have those missing ranks fulfilled, all reads will be considered in the resultant taxonomic profiles (Figure 7B). Even those reads that had taxa of lower rank levels assigned (e.g. Cellular organisms, Bacteria) can be considered in profiles of higher rank levels through nodes of type 3 created by the taxallnomy algorithm (e.g. “spKin_in_Cellular organisms”, “Kin_in_Bacteria”).

**Figure 7:**
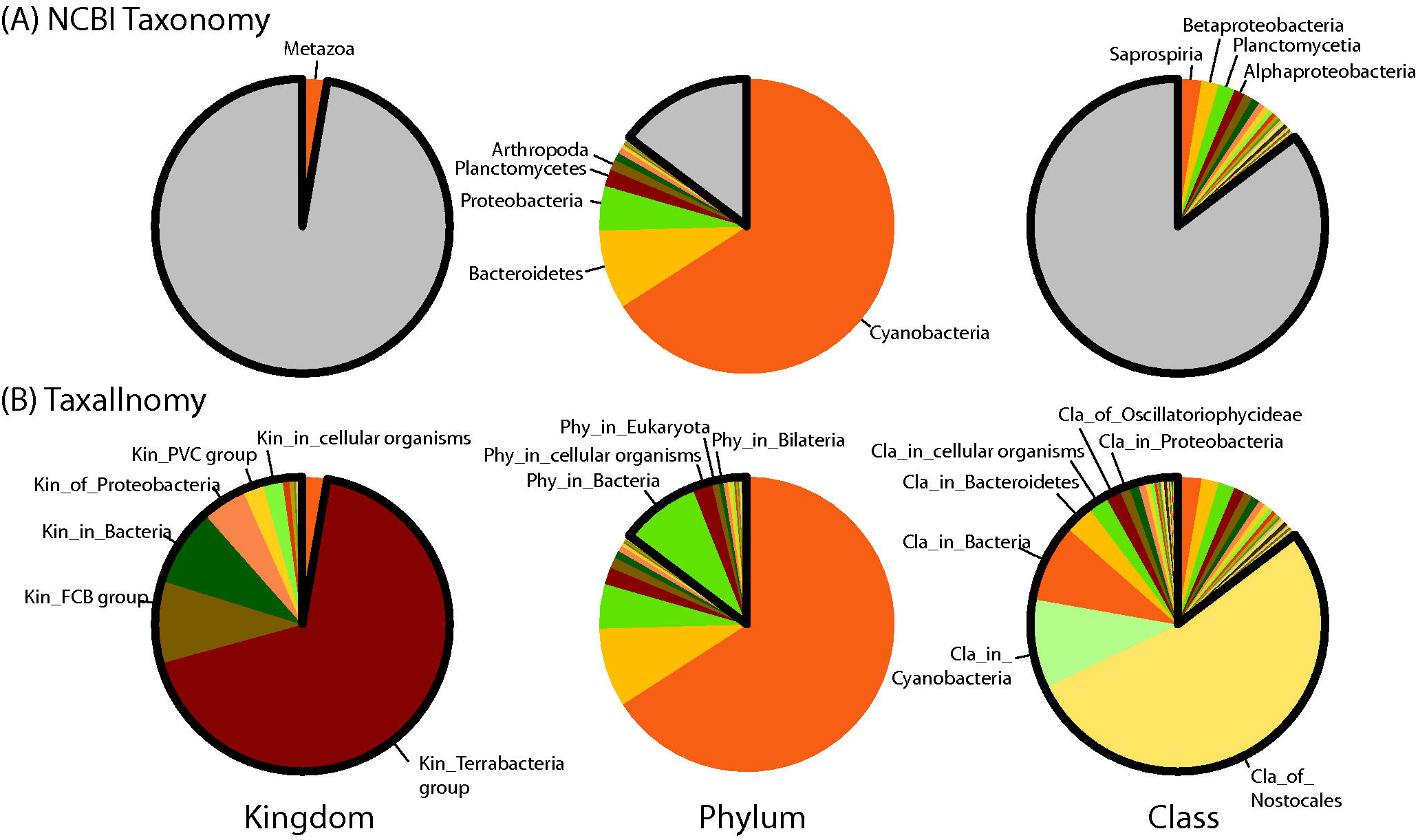
Taxonomic profiles of the metagenomics sample from a tropical freshwater reservoir in Brazil (projectID on MG-RAST:mgp13799) at Kingdom, Phylum and Class ranks, generated by using taxonomic sources from (A) NCBI Taxonomy and (B) taxallnomy databases. For the taxonomic profiles obtained using NCBI Taxonomy, the gray portion of the chart comprises all sequences annotated to taxa that are “NULL” for those ranks in their lineages. These sequences can be further annotated using the lineages retrieved from the taxallnomy database (in bold).

## DISCUSSION

Taxonomy has an extensive history that begins from Aristotle (for review, see (24–27)) and, since then, several approaches have been proposed to classify and name biodiversity. In general, taxonomic databases have two fundamental functions: (1) provide an efficient system of storage and retrieval of taxonomic data; and (2) provide the evolutionary and diversity scenario of the organisms (28,29). To meet one or both functions, two approaches prevail in the current taxonomic databases: (1) the rank-based, which groups organisms in categories of Linnean system (Kingdom, Phylum, Class, etc.); and (2) clade-based, which names monophyletic clades of a phylogenetic tree. Since both classification systems meet very well one of the functions of taxonomy described above (rank-based approach is more practical and clade-based approach is more explanatory) (29), the use of either methodology is a theme under great debate among taxonomists (30–32). Last updates on some databases trends to adopt a clade-based approach (33,34), but other attempts to conciliate both approaches, i.e. NCBI Taxonomy, indicating that taxonomic ranks are still important references.

Taxonomic information provided by NCBI Taxonomy (8) is a valuable resource in several bioinformatics fields. Several tools and software, which have this database as the main subject, have been developed so far either to assist its data retrieval (35–37) or to improve the hierarchical structure by correcting misclassified organism (38,39) or by disambiguating taxonomic names for text mining (40–43). Although the NCBI Taxonomy database has a long life span (since 1991), several reports document challenges presented by the lack of a complete hierarchical rank classification (38,40,44–47). This motivated us to develop taxallnomy, a database that provides a completely hierarchical NCBI Taxonomy rank classification. It is important to emphasize that this work is not meant to propose a new systematic approach for taxonomic classification, but an extension of the broadly used NCBI Taxonomy to facilitate the computational use of the rank-based classification on some bioinformatics approaches.

Since the taxallnomy algorithm could create new nodes (nodes of type 2 and 3) or assign a rank to a preexisting one (nodes of type 1) along with the hierarchical structure, another task performed by the algorithm is to create adequate names to those nodes. Besides the existence of a nomenclature rule to name taxa of a given rank, it would be a complex task to adopt them, since different taxonomic groups have different nomenclature rules (48–50). So, we established generic rules which take advantage of preexisting names and allow easy identification of the rank and the modifications performed by the algorithm in the hierarchical structure.

There is no comprehensive method to address the problem of the lack of a complete hierarchical rank classification, but some solutions have been proposed. The most common and simplest one is the elimination of the “no rank” taxa throughout the lineage (38,40,45). More sophisticated solutions fill the missing ranks by taking the taxon name of the first taxon of a lower (47) or of a higher rank level (46). These solutions are similar to the procedures used by the taxallnomy algorithm to create the nodes of type 2 and 3. Type 2 nodes are created whenever there is no node between two taxa of non-consecutive ranks and take the name of the first ranked taxon of the higher rank level as in (46). The preference to take the name of a higher rank level taxon instead of the lower one is conceptual. For instance, if we have a node “X” of a Phylum rank that has two child nodes “Y” and “Z”, both of the Superclass rank, the Subphylum rank is missing on both Y and Z lineages. By taking the name of the node of lower rank level (node X) to name the missing rank, both Y and Z nodes would have the same Subphylum (“sbPhy of X”). On the other hand, by taking the node of higher rank level (nodes “Y” and “Z”), both nodes will be in distinct Subphylums (“sbPhy of Y” and “sbPhy of Z”). In theory, we do not know if those lineages are actually of the same Subphylum, so, it would be preferable to separate them in different Subphylums instead of putting them in the same group. The node of type 3, on the other hand, is created whenever a lineage lacks higher rank levels. The algorithm takes the name of the last node of the lineage similarly to (47) since there is no other reasonable taxon in which we could take advantage to name the new node.

Besides the creation and nomination of new nodes based on ranked taxa, a remarkable feature performed by the taxallnomy algorithm is the rank assignment of a “no rank” taxon. Taxa with “no rank” status are spread throughout the tree and usually are discarded by users or software that require a hierarchically complete tree. In this work, we show that we could take advantage of “no rank” taxa to fulfill lineages with missing ranks and assist in generating a completely hierarchical taxonomic tree. It is worth mentioning that keeping the “no rank” taxa in the final tree as many as possible is important to preserve the groups already structured by the taxonomic tree. Thus, the current algorithm performs the rank assignment procedure to assign ranks to as many “no rank” taxa as possible. By this, the algorithm has assigned a rank to more than 99% of all “no rank” taxa without disarranging the rank hierarchy already established by the ranked taxa. In the algorithm, we also established a priority scale among the ranks (Table 1) to help in choosing a single rank to be assigned to a “no rank” node with two or more candidate ranks. This procedure favors those most frequent ranks to be selected to assign a “no rank” node (nodes of type 1). We did not note a published report that takes advantage of “no rank” taxa to fulfill all missing ranks. However, a similar but simpler approach can be found in the function “reformat” of a tool named TaxonKit (37), which could address this problem in some bioinformatics applications.

## CONCLUSION

Several bioinformatics analyses and tools rely on the taxonomic information provided by NCBI Taxonomy. However, working with or querying data by taxonomic rank is not trivial because of the absence of some ranks in the taxonomic lineages and the presence of taxa without a rank throughout the taxonomic tree. In this work, we address this issue by developing an algorithm that takes the taxonomic tree from NCBI Taxonomy and makes it hierarchically complete according to the taxonomic ranks. The final tree was named taxallnomy and it has 33 hierarchical levels correspondent to the 33 taxonomic ranks that comprise the NCBI Taxonomy. From the taxallnomy database, the user can retrieve the complete taxonomic lineage with 33 nodes, all of them with a taxonomic rank, to all taxa available in the NCBI Taxonomy. Taxallnomy is applicable to any bioinformatics analyses that depend on the information from NCBI Taxonomy. Here, we demonstrated its application by embedding taxonomic information of a specified rank to a BLAST result and phylogenetic tree; and by making metagenomics profiles. All resources of taxallnomy database is available at bioinfo.icb.ufmg.br/taxallnomy.

## ACKNOWLEDGMENT

We are grateful to Dr. Darren Natale from Protein Information Resource (PIR) for his valuable suggestions to improve our work; to Ph.D. Marcele Laux from Universidade Federal de Minas Gerais (UFMG) for lending us samples and support on metagenomics analysis showed in this work; and to Msc. Edgar Lacerda de Aguiar from Centro Federal de Educação Tecnológica de Minas Gerais (CEFET-MG), which tested and contributed with several suggestions to improve the taxallnomy web interface.

Data available at: http://bioinfo.icb.ufmg.br/taxallnomy

## REFERENCES

1. Roskov, Y., Abucay, L., Orrell, T., et al. (2016) Species 2000 & ITIS Catalogue of Life. Species 2000 & ITIS Catalogue of Life http://www.catalogueoflife.org/ (accessed Jul 8, 2016).

2. Maddison, D. R. and Schulz, K.-S. The Tree of Life Project. The Tree of Life Project http://tolweb.org (accessed Feb 20, 2017).

3. Parr, C. S., Wilson, N., Leary, P., et al. (2014) The Encyclopedia of Life v2: Providing Global Access to Knowledge About Life on Earth. Biodivers. Data J., 2, e1079.

4. GBIF.org (2019) GBIF Home Page. GBIF Home Page https://www.gbif.org/ (accessed Nov 5, 2019).

5. Froese, R. and Pauly, D. (2019) FishBase. FishBase http://www.fishbase.org (accessed May 18, 2020).

6. AmphibiaWeb. AmphibiaWeb https://amphibiaweb.org (accessed May 18, 2020).

7. AnimalBase Project Group (2005) AnimalBase. Early zoological literature online. AnimalBase. Early zoological literature online http://www.animalbase.uni-goettingen.de (accessed May 18, 2020).

8. Federhen, S. (2012) The NCBI Taxonomy database. Nucleic Acids Res., 40, D136–D143.

9. Cochrane, G., Karsch-Mizrachi, I., Takagi, T., et al. (2016) The International Nucleotide Sequence Database Collaboration. Nucleic Acids Res., 44, D48–D50.

10. Consortium, T. U. (2015) UniProt: a hub for protein information. Nucleic Acids Res., 43, D204–D212.

11. Aken, B. L., Ayling, S., Barrell, D., et al. (2016) The Ensembl gene annotation system. Database, 2016.

12. Finn, R. D., Bateman, A., Clements, J., et al. (2014) Pfam: the protein families database. Nucleic Acids Res., 42, D222–230.

13. Schultz, J., Milpetz, F., Bork, P., et al. (1998) SMART, a simple modular architecture research tool: identification of signaling domains. Proc. Natl. Acad. Sci. U. S. A., 95, 5857–5864.

14. Mi, H., Muruganujan, A. and Thomas, P. D. (2013) PANTHER in 2013: modeling the evolution of gene function, and other gene attributes, in the context of phylogenetic trees. Nucleic Acids Res., 41, D377–D386.

15. Altenhoff, A. M., Škunca, N., Glover, N., et al. (2015) The OMA orthology database in 2015: function predictions, better plant support, synteny view and other improvements. Nucleic Acids Res., 43, D240–D249.

16. Kozomara, A. and Griffiths-Jones, S. (2011) miRBase: integrating microRNA annotation and deep-sequencing data. Nucleic Acids Res., 39, D152–D157.

17. Berman, H. M., Westbrook, J., Feng, Z., et al. (2000) The Protein Data Bank. Nucleic Acids Res., 28, 235–242.

18. Kolesnikov, N., Hastings, E., Keays, M., et al. (2015) ArrayExpress update—simplifying data submissions. Nucleic Acids Res., 43, D1113–D1116.

19. Kanehisa, M., Furumichi, M., Tanabe, M., et al. (2017) KEGG: new perspectives on genomes, pathways, diseases and drugs. Nucleic Acids Res., 45, D353–D361.

20. Altschul, S. F., Gish, W., Miller, W., et al. (1990) Basic local alignment search tool. J. Mol. Biol., 215, 403–410.

21. Huson, D. H., Auch, A. F., Qi, J., et al. (2007) MEGAN analysis of metagenomic data. Genome Res., 17, 377–386.

22. Keegan, K. P., Glass, E. M. and Meyer, F. (2016) MG-RAST, a Metagenomics Service for Analysis of Microbial Community Structure and Function. Methods Mol. Biol. Clifton NJ, 1399, 207–233.

23. Mitchell, A. L., Scheremetjew, M., Denise, H., et al. (2018) EBI Metagenomics in 2017: enriching the analysis of microbial communities, from sequence reads to assemblies. Nucleic Acids Res., 46, D726–D735.

24. Mishler, B. D. (2009) Three centuries of paradigm changes in biological classification: Is the end in sight? TAXON, 58, 61–67.

25. Raven, P. H., Berlin, B. and Breedlove, D. E. (1971) The origins of taxonomy. Science, 174, 1210–1213.

26. Mayr, E. (1982) The growth of biological thought: Diversity, evolution, and inheritance. The growth of biological thought: Diversity, evolution, and inheritance; Harvard University Press, (1982).

27. Stevens, P. F. (1994) The development of biological systematics: Antoine-Laurent de Jussieu, nature, and the natural system. The development of biological systematics: Antoine-Laurent de Jussieu, nature, and the natural system; Columbia University Press, (1994).

28. Mayr, E. and Bock, W. J. (2002) Classifications and other ordering systems. J. Zool. Syst. Evol. Res., 40, 169–194.

29. Dubois, A. (2007) Phylogeny, taxonomy and nomenclature:the problem of taxonomic categories and of nomenclatural ranks. Zootaxa, 1519, 27–68.

30. Nixon, K. C., Carpenter, J. M. and Stevenson, D. W. (2003) The PhyloCode is fatally flawed, and the “Linnaean” System can easily be fixed. Bot. Rev., 69, 111.

31. Rieppel, O. (2006) The PhyloCode: a critical discussion of its theoretical foundation. Cladistics, 22, 186–197.

32. Pennisi, E. (2001) Linnaeus’s Last Stand? Science, 291, 2304–2307.

33. Adl, S. M., Bass, D., Lane, C. E., et al. (2019) Revisions to the Classification, Nomenclature, and Diversity of Eukaryotes. J. Eukaryot. Microbiol., 66, 4–119.

34. Chase, M. W., Christenhusz, M. J. M., Fay, M. F., et al. (2016) An update of the Angiosperm Phylogeny Group classification for the orders and families of flowering plants: APG IV. Bot. J. Linn. Soc., 181, 1–20.

35. Stajich, J. E., Block, D., Boulez, K., et al. (2002) The Bioperl toolkit: Perl modules for the life sciences. Genome Res., 12, 1611–1618.

36. Huerta-Cepas, J., Serra, F. and Bork, P. (2016) ETE 3: Reconstruction, Analysis, and Visualization of Phylogenomic Data. Mol. Biol. Evol., 33, 1635–1638.

37. Shen, W. and Xiong, J. (2019) TaxonKit: a cross-platform and efficient NCBI taxonomy toolkit. bioRxiv, 513523.

38. McDonald, D., Price, M. N., Goodrich, J., et al. (2012) An improved Greengenes taxonomy with explicit ranks for ecological and evolutionary analyses of bacteria and archaea. ISME J., 6, 610–618.

39. Kozlov, A. M., Zhang, J., Yilmaz, P., et al. (2016) Phylogeny-aware identification and correction of taxonomically mislabeled sequences. Nucleic Acids Res., 44, 5022–5033.

40. Naderi, N., Kappler, T., Baker, C. J. O., et al. (2011) OrganismTagger: detection, normalization and grounding of organism entities in biomedical documents. Bioinforma. Oxf. Engl., 27, 2721–2729.

41. Wei, C.-H., Kao, H.-Y. and Lu, Z. (2012) SR4GN: A Species Recognition Software Tool for Gene Normalization. PLOS ONE, 7, e38460.

42. Pafilis, E., Frankild, S. P., Fanini, L., et al. (2013) The SPECIES and ORGANISMS Resources for Fast and Accurate Identification of Taxonomic Names in Text. PLOS ONE, 8, e65390.

43. Boyle, B., Hopkins, N., Lu, Z., et al. (2013) The taxonomic name resolution service: an online tool for automated standardization of plant names. BMC Bioinformatics, 14, 16.

44. Porter, T. M., Gibson, J. F., Shokralla, S., et al. (2014) Rapid and accurate taxonomic classification of insect (class Insecta) cytochrome c oxidase subunit 1 (COI) DNA barcode sequences using a naïve Bayesian classifier. Mol. Ecol. Resour., 14, 929–942.

45. Ekstrom, A. and Yin, Y. (2016) ORFanFinder: automated identification of taxonomically restricted orphan genes. Bioinformatics, 32, 2053–2055.

46. García-López, R., Vázquez-Castellanos, J. F. and Moya, A. (2015) Fragmentation and Coverage Variation in Viral Metagenome Assemblies, and Their Effect in Diversity Calculations. Front. Bioeng. Biotechnol., 3, 141.

47. Guillou, L., Bachar, D., Audic, S., et al. (2013) The Protist Ribosomal Reference database (PR2): a catalog of unicellular eukaryote Small Sub-Unit rRNA sequences with curated taxonomy. Nucleic Acids Res., 41, D597–D604.

48. ICZN (International Commission on Zoological Nomenclature) (1999) International code of zoological nomenclature; Fourth edition; International Trust for Zoological Nomenclature, (1999).

49. (1992) International Code of Nomenclature of Bacteria: Bacteriological Code, 1990 Revision. International Code of Nomenclature of Bacteria: Bacteriological Code, 1990 Revision; Lapage, S. P., Sneath, P. H. A., Lessel, E. F., et al. (eds.); ASM Press, Washington (DC), (1992).

50. (2018) International Code of Nomenclature for algae, fungi, and plants. International Code of Nomenclature for algae, fungi, and plants; Turland, N., Wiersema, J., Barrie, F., et al. (eds.); Regnum Vegetabile; Koeltz Botanical Books, (2018); Vol. 159.

